# Astrocytic Gi-GPCR activation enhances stimulus-evoked extracellular glutamate

**DOI:** 10.1101/2022.05.12.491656

**Authors:** Trisha V. Vaidyanathan, Vincent Tse, Esther M. Lim, Kira E. Poskanzer

## Abstract

Astrocytes perform critical functions in the nervous system, many of which are dependent on neurotransmitter-sensing through G protein-coupled receptors (GPCRs). However, whether specific astrocytic outputs follow specific GPCR activity remains unclear, and exploring this question is critical for understanding how astrocytes ultimately influence brain function and behavior. We previously showed that astrocytic Gi-GPCR activation is sufficient to increase slow-wave neural activity (SWA) during sleep when activated in cortical astrocytes^1^. Here, we investigate the outputs of astrocytic Gi-GPCRs, focusing on the regulation of extracellular glutamate and GABA, by combining *in vivo* fiber photometry recordings of the extracellular indicators iGluSnFR and iGABASnFR with astrocyte-specific chemogenetic Gi-GPCR activation. We find that Gi-GPCR activation does not change spontaneous dynamics of extracellular glutamate or GABA. However, Gi-GPCR activation does specifically increase visual stimulus-evoked extracellular glutamate. Together, these data point towards a complex relationship between astrocytic inputs and outputs *in vivo* that may depend on behavioral context. Further, they suggest an extracellular glutamate-specific mechanism underlying some astrocytic Gi-GPCR-dependent behaviors, including the regulation of sleep SWA.

## Introduction

Astrocytes and neurons are involved in a constant conversation via extracellular molecules. Astrocytes sense neurotransmitter signals via metabotropic G protein-coupled receptors (GPCRs)^2^, which results in astrocyte activation via elevations in calcium (Ca^2+^)^3,4^. In turn, astrocytes can regulate neuronal activity and behavior^1,5–10^. However, these are general outlines of this astrocyte-neuron cross-talk. Whether particular astrocytic outputs can be mapped back to specific upstream astrocytic GPCR activity remains largely undetermined.

Astrocytes in the cerebral cortex are well suited to regulate neuronal network activity via their anatomical interconnectedness and ability to bidirectionally communicate with large populations of neurons^11^. In fact, cortical astrocytic GPCR signaling has been shown to be critical for the regulation of non-rapid eye movement (NREM) sleep^1,12,13^, a behavior critical for learning and memory involving the rhythmic activity of large populations of cortical neurons. Previously, we showed a family of GPCRs coupled to the Gi alpha subunit (Gi-GPCRs) in cortical astrocytes is sufficient to regulate NREM slow-wave activity (SWA), a marker of sleep depth^1^. Further, Gi-GPCR activation of astrocytes in other brain regions has been recently shown to affect other behaviors during wake^6,7,14,15^. While Gi-GPCR activation in astrocytes increases intracellular Ca^2+^ signaling^1,7,9,16^, it remains unknown which downstream astrocytic outputs drive the observed changes in SWA.

The cortex plays a critical role in the generation of SWA^17–25^, which can be explained mechanistically by actions of both the excitatory neurotransmitter glutamate^18,26^ and the inhibitory neurotransmitter GABA^27–30^. Astrocytes respond to both glutamate and GABA through Gi-GPCRs (mGluR3 and GABA_B_ receptors, respectively, in adults)^7,16,31^, and regulate extracellular glutamate ([glu]_e_) and extracellular GABA ([GABA]_e_) by uptake from the extracellular space via transporters^32^. Thus, the regulation of [glu]_e_ and [GABA]_e_ are attractive candidates as 1) functional outputs of Gi-GPCR activation and 2) potential mechanisms of astrocytic regulation of sleep SWA.

Here, we combine astrocyte-specific chemogenetic manipulations of the Gi-GPCR pathway with fiber photometry recordings of the extracellular neurotransmitter indicators iGluSnFR and iGABASnFR2^33^ to investigate how cortical astrocyte Gi-GPCR activation affects [glu]_e_ and [GABA]_e_ *in vivo*. We find that Gi-GPCR activation does not change baseline spontaneous [glu]_e_ or [GABA]_e_ dynamics. However, using a light flash visual stimulus, we found that astrocytic Gi-GPCR activation did increase the amplitude specifically of evoked [glu]_e_, but not [GABA]_e_ in primary visual cortex (V1). Together, these data point toward a specific glutamatergic output of astrocytic Gi-GPCR activation in one context, and suggests that control of [glu]_e_ may be a mechanism for other Gi-GPCR-dependent astrocytic functions, including sleep regulation.

## Results

### Recording extracellular glutamate and GABA in freely moving mice

To investigate the functional output of astrocytic Gi-GPCR signaling *in vivo*, we combined astrocytic Gi-GPCR activation with recordings of [glu]_e_ and [GABA]_e_ in freely moving mice (**Fig. 1a**). We focused on V1, where we previously showed that Gi-GPCR activation in astrocytes increases intracellular Ca^2+^ activity and regulates SWA^1^. Prior to experiments, mice were co-injected with two viruses (**Fig. 1b**). To activate the Gi pathway in astrocytes, we used Designer Receptors Exclusively Activated by Designer Drugs (DREADDs)^34^, injecting mice with *AAV-GFAP-hM4Di-mCherry* to express the human M4 muscarinic receptor DREAD (hM4Di) in V1 astrocytes. The second virus injected was either *AAV-GFAP-SF-iGluSnFR*.*A184S*^35^ or a updated variant of *AAV-GFAP-iGABASnFR2*^36^ (GENIE Project team, Janelia Research Campus, HHMI), to express a fluorescent extracellular indicator for glutamate or GABA, respectively, in astrocytes. Immunohistochemistry confirmed astrocytic expression of all viruses (**Fig. 1b**, NeuN co-localization for iGluSnFR: 0.59 ± 0.98%, iGABASnFR2: 0.40 ± 0.31%, Gi-DREADD: 13.64 ± 5.47%). After waiting 2–3 weeks for recovery and expression, we performed fiber photometry recordings of iGluSnFR or iGABASnFR2 as mice moved around a circular chamber (**Fig. 1a, c–d**). We examined the effect of Gi-activation on [glu]_e_ or [GABA]_e_ via I.P. administration of the hM4Di agonist clozapine-N-oxide (CNO, 1mg/kg), compared to control I.P. injections of saline. In our previous study^1^, astrocyte Gi-GPCR-induced Ca^2+^ dynamics were precisely characterized on a subcellular scale using *in vivo* two-photon imaging. Here, we sought to capture how [glu]_e_ and [GABA]_e_ dynamics changed across large cellular populations in response to population-wide astrocytic activation. We observed robust, spontaneous fluctuations in both the iGluSnFR and iGABASnFR2 signal as mice moved around the circular chamber (**Fig. 1c–d**). In addition to spontaneous dynamics, we also elicited [glu]_e_ and [GABA]_e_ events using visual stimuli in the form of a blue LED flash (**Fig. 1a**). We observed consistent increases to the light flash when recording either iGluSnFR or iGABASnFR2 (**Fig. 1e–g**).

**Figure 1:**
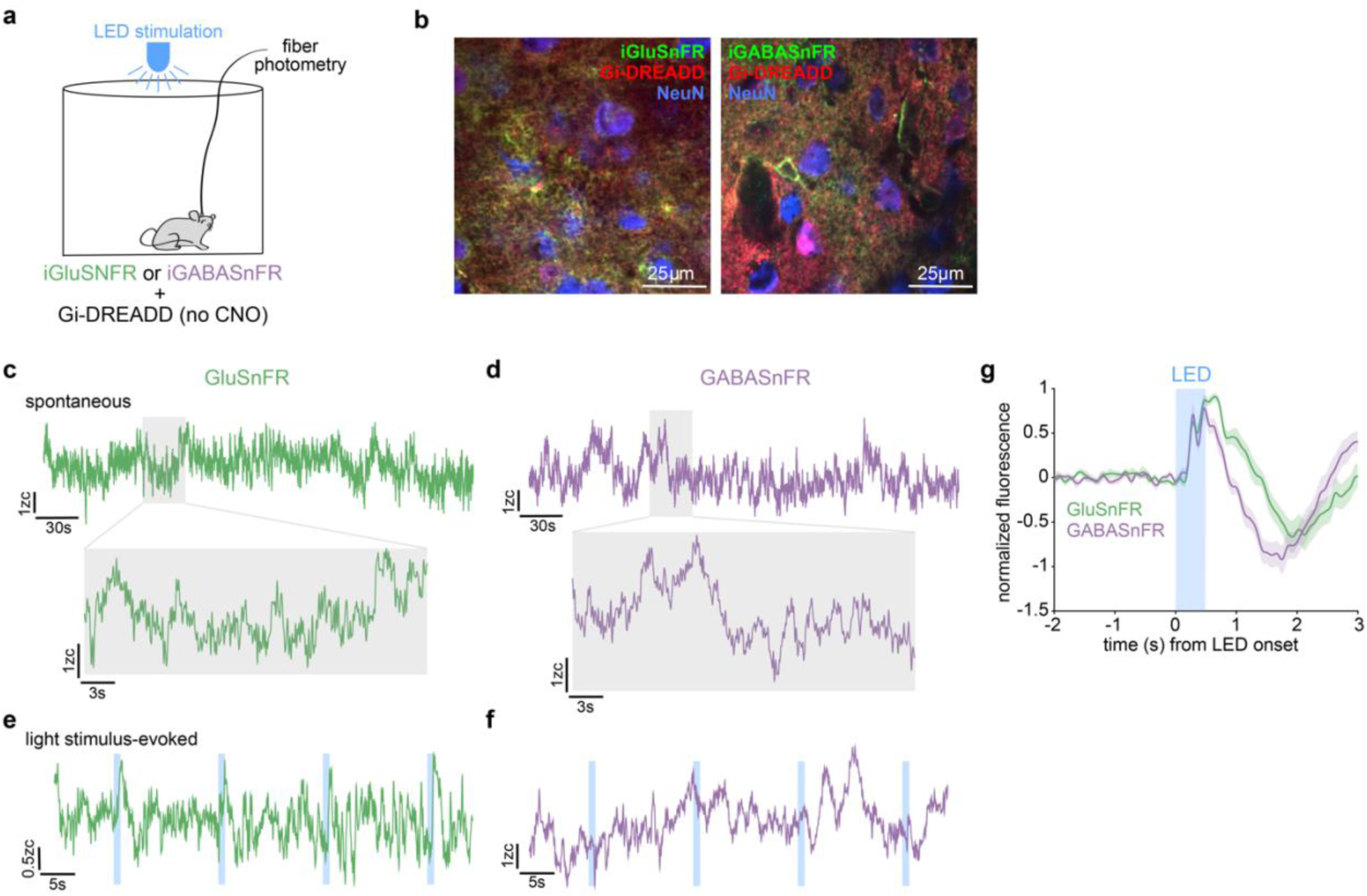
*In vivo* recording spontaneous and stimulus-evoked extracellular glutamate and GABA in freely moving mice. **(a)** Experimental *in vivo* fiber photometry setup. Mice were co-injected with either *GFAP-iGluSnFR* or *GFAP-iGABASnFR2* and *GFAP-hM4D(Gi)-mCherry* AAVs. Extracellular glutamate and GABA were recorded using fiber photometry as mice moved freely in a circular chamber. **(b)** Representative immunohistochemistry images showing expression of Gi-DREADDs (red), NeuN (blue), and iGluSnFR (left, green) or iGABASnFR2 (right, green). **(c–d)** Example of spontaneous iGluSnFR (c) dynamics over 5min (top) and 30s (bottom) and spontaneous iGABASnFR2 (d) dynamics over 5min (top) and 30s (bottom). **(e–f)** Example of consistent iGluSnFR (e) and iGABASnFR2 (f) responses to light flashes (blue bars) **(g)** Light-evoked average photometry response with iGluSnFR (green) and iGABASnFR2 (purple) (iGluSnFR: n = 10 mice; iGABASnFR2: n = 9 mice. 78 stimuli per mouse. Error bars = SEM).

### Gi-DREADD activation of cortical astrocytes does not change spontaneous extracellular glutamate or GABA dynamics

To investigate how intracellular astrocytic Gi-GPCR signaling affects [glu]_e_ and [GABA]_e_ in V1, we set out to measure spontaneous [glu]_e_ and [GABA]_e_ activity with fiber photometry in mice expressing astrocytic Gi-DREADDs, comparing CNO-injected animals to saline-injected controls (**Fig. 2a**). As a control, we first confirmed that this Gi-DREADD manipulation resulted in calcium (Ca^2+^) increases with the same CNO dose (1mg/kg), as we previously published^1^. To do so, we injected a separate cohort of mice with viruses to co-express astrocytic GCaMP and Gi-DREADDs and performed fiber photometry recordings of Ca^2+^ activity following either CNO or saline injections (**Figure 2–figure supplement 1a– b**). Post-recording immunohistochemistry confirmed astrocytic expression of GCaMP (1.86 ± 1.35% NeuN co-localization) and Gi-DREADDs (5.79 ± 1.45% NeuN co-localization) (**Figure 2–figure supplement 1b**). As in our previous study, CNO administration increased Ca^2+^ event frequency (**Figure 2–figure supplement 1d-e**) but did not change the magnitude of Ca^2+^ events (**Figure 2–figure supplement 1f**).

**Figure 2:**
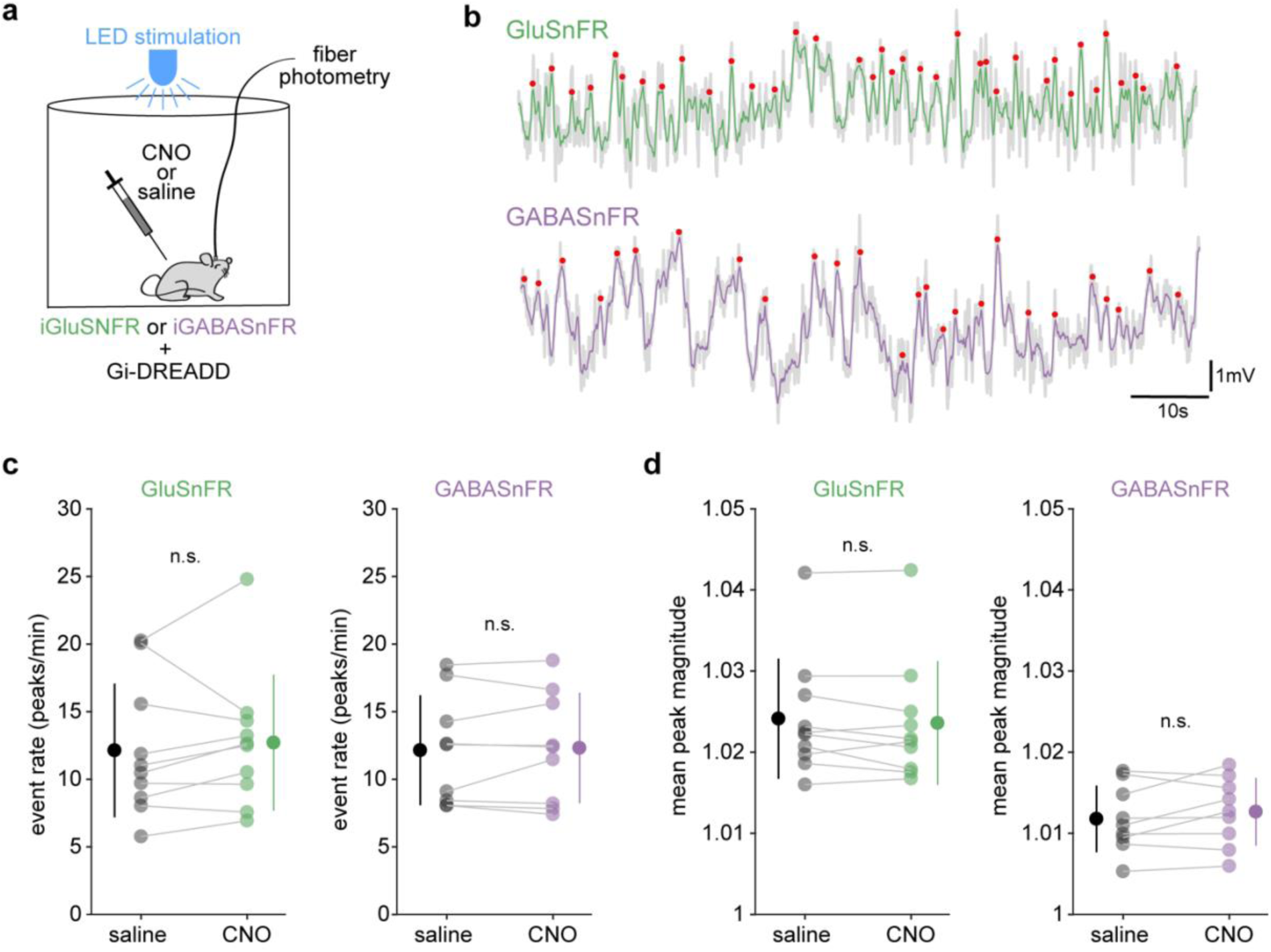
Astrocyte Gi-DREADD activation does not change spontaneous iGluSnFR or iGABASnFR2 dynamics. **(a)** Experimental paradigm. Photometry recordings were performed after I.P. injection of either CNO or saline, including 30min of spontaneous activity. **(b)** Example traces of spontaneous iGluSnFR (top, green) and iGABASnFR2 (bottom, purple) dynamics. Events were identified by automatically detecting peaks (red dots). **(c)** Event rate (peaks/min) was the same after saline and CNO administration for both iGluSnFR (left) and iGABASnFR2 (right) recordings. **(d)** Spontaneous event magnitude was equivalent after saline and CNO administration for both iGluSnFR (left) and iGABASnFR2 (right) recordings. (For panels c–d: iGluSnFR, n = 9 mice, iGABASnFR2, n = 9 mice. 78 stimuli per mouse, paired t-test, error bars = SD).

We next measured iGluSnFR and iGABASnFR2 after Gi-DREADD activation. To do this, we quantified transients by automatically detecting local maxima (**Fig. 2b, see Methods**). In so doing, we found that Gi-GPCR activation did not change the frequency (**Fig. 2c**), nor the amplitude (**Fig. 2d**) of iGluSnFR or iGABASnFR2 transients. These results suggest that Gi-GPCR signaling in astrocytes does not significantly contribute to basal levels of glutamatergic or GABAergic signaling in V1 circuits, at levels detectable via this experimental paradigm. Since glutamate and GABA are under tight regulation^37^, it may indeed be expected that activation of astrocytes does not result in large changes to endogenous [glu]_e_ or [GABA]_e_ dynamics. Thus, we next investigated whether astrocytic Gi-GPCR activation changes [glu]_e_ or [GABA]_e_ in a stimulus-evoked experimental paradigm.

### Gi-DREADD activation does not change the decay rate of stimulus-evoked extracellular glutamate or GABA

Astrocytes express transporters for glutamate (GLT-1, GLAST)^38,39^ and GABA (GAT-1, GAT-3)^40,41^ at high levels. These transporters are a primary mechanism for neurotransmitter uptake in the CNS and astrocytic uptake of glutamate and GABA is critical for the finely tuned balance of excitation and inhibition in the cortex^32,38,42–44^. We hypothesized here that Gi-GPCR activation may regulate glutamate or GABA uptake. To test this, we examined whether Gi-DREADD activation via CNO would change the decay rate—a measure of neurotransmitter uptake^42,43^—of iGluSnFR or iGABASnFR2 events.

To obtain a reliable measure of decay, we used a visual stimulus to elicit iGluSnFR and iGABASnFR2 events in V1 (**Fig. 3a–b**, see Methods). LED flashes resulted in consistent, stereotyped responses of both iGluSnFR and iGABASnFR2: a rapid increase in fluorescence with a brief lag from LED onset (132 ± 64ms for iGluSnFR, 112 ± 74ms for iGABASnFR2), followed by a decrease in fluorescence that dipped below baseline levels before returning to baseline (**Fig. 3b**). In these responses, we observed two peaks (“early” and “late”) for both iGluSnFR and iGABASnFR2 before the fluorescence decreased to baseline (**Fig. 3c, e**). An early and late response to a light flash has been reported previously in mice, cats, and humans^45–48^. Indeed, the late component has been reported to occur >300ms and ≤1s following a light flash^47,48^, similar to the latency we observed (433 ± 46ms for iGluSnFR and 433 ± 16ms for iGABASnFR2).

**Figure 3:**
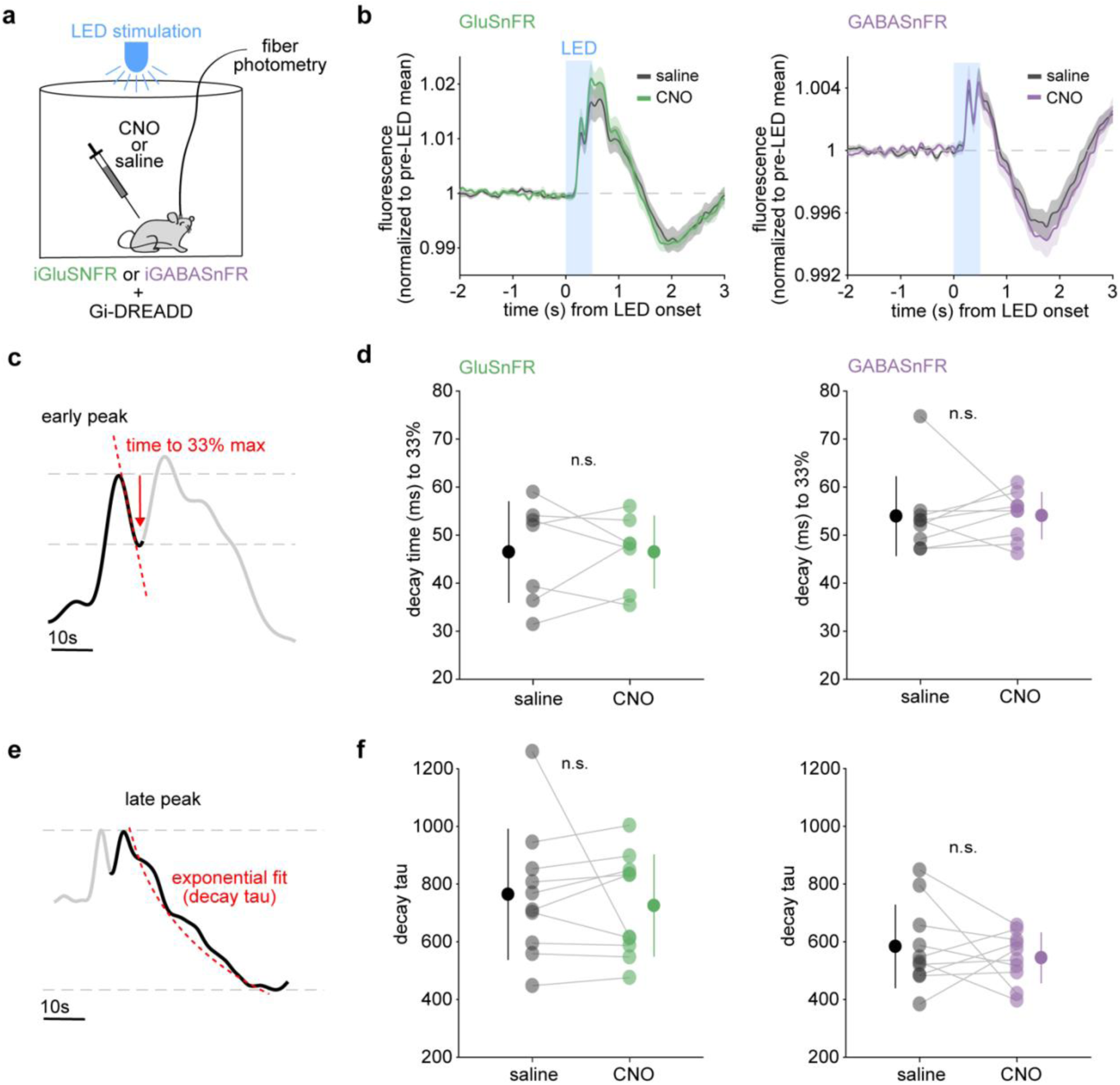
Gi-DREADD activation does not change the decay time of stimulus-evoked iGluSnFR or iGABASnFR2. **(a**) Experimental *in vivo* fiber photometry setup. Mice were co-injected with either *GFAP-iGluSnFR* or *GFAP-iGABASnFR2* and *GFAP-hM4D(Gi)-mCherry* AAVs. After I.P. injection of either 1mg/kg CNO or saline, extracellular glutamate and GABA were recorded using fiber photometry as mice moved freely in a circular chamber with intermittent LED stimuli. **(b)** Light-evoked average iGluSnFR (left) and iGABASnFR2 (right) response is similar after CNO or saline injection. (iGluSnFR: n = 10 mice; iGABASnFR2: n = 9 mice. For all, 78 stimuli per mouse. Error bars = SEM). **(c)** Light-evoked responses had two peaks, an “early” and a “late” response. The response decay for the early peak was quantified by calculating the time (ms) for the first peak response to decay to 33% of the max. **(d)** The early light-evoked response for iGluSnFR (left) and iGABASnFR2 (right) had similar decay times after CNO and saline administration (iGluSnFR: n = 7 mice; iGABASnFR2: n = 9 mice. For all, 78 stimuli per mouse, paired t-test, error bars = SD). **(e)** The decay for the late peak was quantified by fitting an exponential function to the mean trace for each mouse. **(f)** The late light-evoked response for iGluSnFR (left) and iGABASnFR2 (right) had similar decay times after CNO and saline administration (iGluSnFR: n = 10 mice; iGABASnFR2: n = 9 mice. For all, 78 stimuli per mouse, paired t-test, error bars = SD).

To investigate the role of Gi-GPCR signaling in glutamate and GABA uptake, we compared the responses between CNO and saline control conditions. We observed no discernable difference in the overall shape of the LED-evoked response of iGluSnFR or iGABASnFR2 (**Fig. 3b**). Next, we quantified the decay for both the early and late peak responses. The early peak decay was linear, so we computed the time for fluorescence to decay to 33% of the maximum response (**Fig. 3c**). We observed no significant difference in this decay time between saline and CNO conditions for either iGluSnFR or iGABASnFR2 recordings (**Fig. 3d**), suggesting that glutamate and GABA uptake for this initial visual response is not affected by astrocytic Gi-GPCR signaling. Next, we examined the decay for the late response. The decrease in fluorescence following the late response was exponential, so we quantified the decay tau by fitting an exponential curve to this portion of the trace (**Fig. 3e, see Methods**). Similar to the early decay, we found no significant difference in the tau of the late response for iGluSnFR or iGABASnFR2 (**Fig. 3f**). Together, these results suggest that astrocytic Gi-GPCR activation does not affect the regulation of glutamate or GABA uptake.

### Gi-DREADD activation increases the amplitude of stimulus-evoked extracellular glutamate, but not extracellular GABA

While we did not observe a significant change in the amplitude of spontaneous iGluSnFR or iGABASnFR2 events (**Fig. 2d**) with astrocytic Gi-DREADD activation, we wondered whether the amplitude of evoked events might change. To test this, we examined the amplitude of the early and late peak responses separately, as described above (**Fig. 4a, c**). While we observed no significant change in amplitude for the early iGABASnFR2 response (**Fig. 4b, right**), there was a trend towards an increase in the iGluSnFR response (**Fig. 4b, left**). When we quantified the late peak amplitude, we found there was a significant increase in the iGluSnFR response amplitude, but not the iGABASnFR2 amplitude (**Fig. 4d**). This result suggests that astrocytic Gi-GPCR signaling can modulate downstream glutamatergic signaling, but not GABAergic signaling in this experimental context. It is possible that stimulus-evoked changes in glutamate bypass homeostatic mechanisms that prevented a similar increase to be observed in the spontaneous dynamics (**Fig. 2d**). Alternatively, this may be a stimulus-specific mechanism. Previous work from our group has shown that astrocyte Gi-DREADD activation is sufficient to increase SWA in sleep^1^, and glutamate has been shown to be sufficient to generate cortical UP states^18,26^, a major component of SWA. Together, this data support the hypothesis that increased [glu]_e_ is a mechanism by which astrocytic Gi-GPCR signaling modulates SWA.

**Figure 4:**
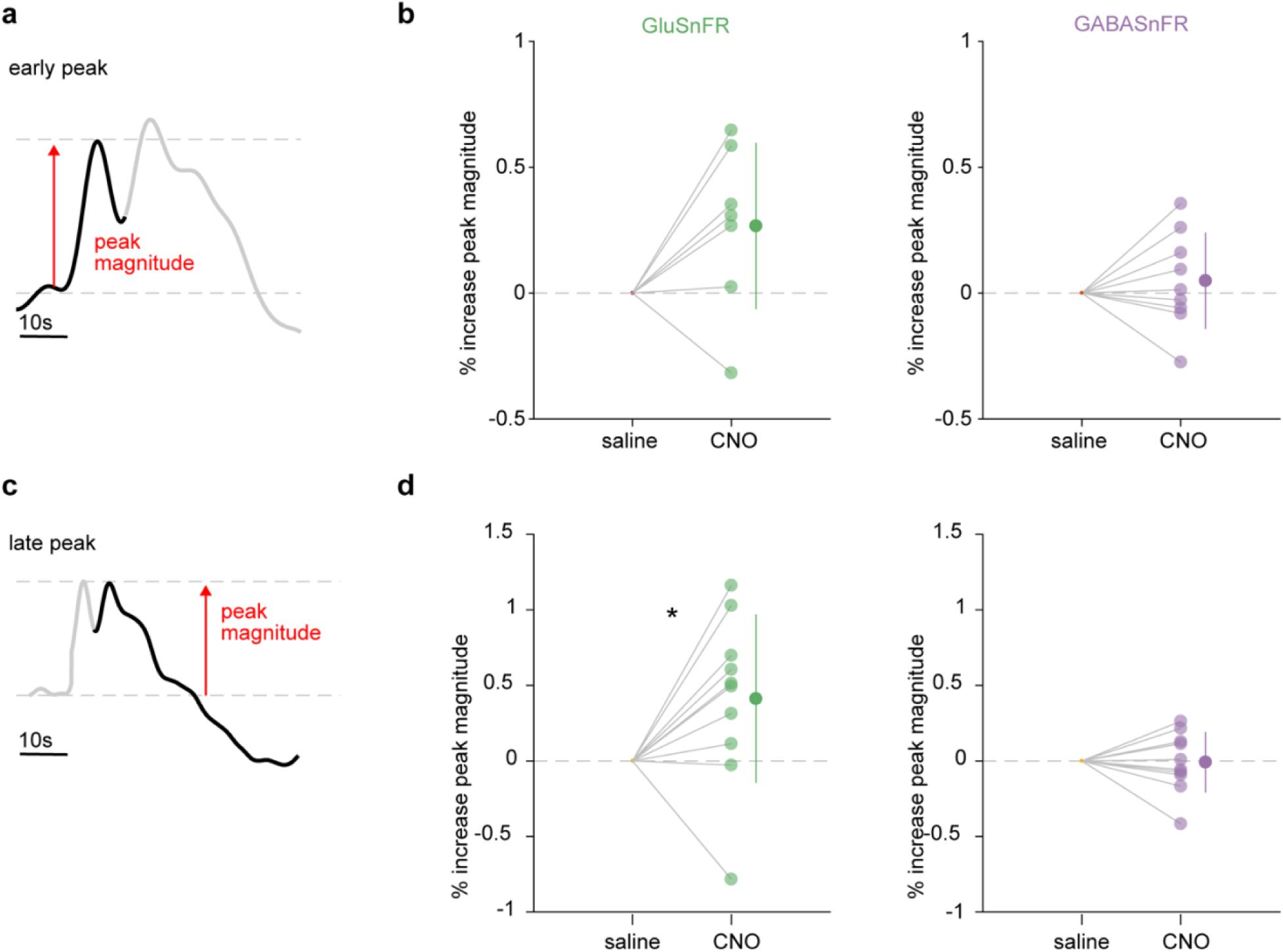
Gi-DREADD activation increases the magnitude of stimulus-evoked iGluSnFR, but not iGABASnFR2. **(a)** The magnitude of the early light-evoked iGluSnFR and iGABASnFR2 response was quantified after I.P. injection of either 1mg/kg CNO or saline. **(b)** The magnitude of the early iGluSnFR light-evoked response trended towards an increase after CNO administration (left), but there was no change for the iGABASnFR2 response. (iGluSnFR: n = 7 mice; iGABASnFR2: n = 9 mice. For all, 78 stimuli per mouse, paired t-test, p = 0.07, error bars = SD). **(c)** The magnitude of the late light-evoked iGluSnFR and iGABASnFR2 response was quantified after CNO or saline administration. **(d)** CNO administration caused a significant increase in the late light-evoked iGluSnFR response (left) but not the late iGABASnFR2 response (right). (iGluSnFR: n = 9 mice; iGABASnFR2: n = 9 mice. For all, 78 stimuli per mouse, paired t-test, error bars = SD).

These data do not allow us to distinguish whether increases in [glu]_e_ are ultimately neuron-or astrocyte-derived, although the chemogenetic activation of Gi-GPCRs is limited to astrocytes. However, previous work has demonstrated that astrocytic Gi-DREADD activation can increase slow-inward currents which are thought to be the result of astrocytic glutamate release^16^. Regardless of cell type, these data point toward a glutamate-specific functional output of astrocytic Gi-GPCR activity.

## Discussion

Using *in vivo* fiber photometry recordings of iGluSnFR and iGABASnFR2 in freely moving mice, we find that astrocyte-specific Gi-GPCR activation does not significantly change spontaneous [glu]_e_ or [GABA]_e_, nor does it change the decay dynamics of evoked [glu]_e_ or [GABA]_e_. However, we do find that Gi-GPCR activation increases the amplitude of evoked [glu]_e_ events, while [GABA]_e_ is unchanged. These data point toward a glutamate-specific output mechanism of Gi-GPCR signaling in specific contexts

### Glutamatergic mechanism of astrocytic SWA regulation

In this study, we focused on the dynamics of the two major excitatory and inhibitory neurotransmitters in response to a type of astrocytic activation which has been used by many research groups in the field in recent years^6,7,14–16^. However, we did not investigate sleep, so do not make any conclusions about SWA mechanisms. Nevertheless, our finding that astrocytic Gi-GPCR activation increased [glu]_e,_ but not [GABA]_e_, events (**Fig. 4d**) may point toward a specific mechanism underlying the previously reported astrocytic control of SWA^1^, through the regulation of cortical UP states. Glutamate is sufficient to initiate cortical UP states^18,26^ while synchronous DOWN states may be driven specifically by inhibitory GABA signaling^17,27,28,30^. Indeed, astrocytes have been linked specifically to UP states *ex vivo* previously^26^.

If the observed increase in [glu]_e_ amplitude is responsible for the Gi-GPCR control of SWA, it would suggest relatively fast (within ms) regulation of [glu]_e_ by astrocytes. Astrocytes dynamically regulate glutamate^49^ and can shape excitatory post-synaptic events on a ms timescale via glutamate uptake^44^. However, we found no change in the iGluSnFR decay time—a measure of glutamate uptake—with Gi-GPCR activation (**Fig. 3d, f left**), suggesting that glutamate uptake is not dependent on the Gi-GPCR pathway. Although contentious^50^, Gi-GPCR activation may instead result in astrocytic release of glutamate. This is supported by previous work showing astrocytic Gi-GPCR activation increases slow-inward currents^16^. Whether changes in [glu]_e_ observed here occur via astrocytic vesicular release^51–54^ or the reversal of glutamate transporters^55^ remain for future research.

### Functional outputs of astrocytic Gi-GPCR signaling

Beyond the regulation of sleep, Gi-GPCR signaling in astrocytes has been linked to behaviors during the wake state^6,7,14,15^, and the data presented here provide important insight into general astrocyte Gi-GPCR signaling. Only recently has chemogenetic astrocyte Gi-GPCR signaling been shown to increase astrocyte Ca^2+^ through an IP_3_R2-dependent pathway^17^, unlike in neurons^56^. The downstream functional outputs that result from activation of this signaling pathway to ultimately affect neuronal activity and behavior has not yet been widely explored. Our finding that chemogenetic astrocyte Gi-GPCR activation can preferentially change [glu]_e_ but not [GABA]_e_, may help elucidate the mechanisms underlying other astrocyte-dependent behaviors.

The observed change in [glu]_e_ upon Gi-GPCR activation may also underlie a homeostatic role of astrocytes in maintaining excitatory/inhibitory (E/I) balance within the cortex. Indeed, astrocytic sensing of GABA via the Gi-GPCR GABA_B_ receptor has been previously implicated in E/I balance^57^ and astrocytes can increase glutamate transmission upon sensing of GABA via GABA_B_ receptors *ex vivo*^16,58^. To identify the specific type of Gi-GPCR responsible for the observed increase in [glu]_e_ here, further work will be necessary, such as loss-of-function experiments to genetically silence GABA_B_ receptors.

### Spontaneous and stimulus-evoked dynamics

We observed a significant change in [glu]_e_ amplitude with Gi-GPCR activation during visual stimulus-evoked [glu]_e_ dynamics, but not during spontaneous activity. This difference underscores the complexity of these signals and highlights the varying role astrocytes may play in circuit function under different behavioral contexts. One explanation for the difference we observe may be that Gi-GPCR activation always increases [glu]_e_, but the increase is masked during spontaneous activity due to compensatory mechanisms that are not present during strong visual stimuli. Indeed, the increase in amplitude observed with visual stimuli was relatively small. Increasing the amount of Gi-GPCR activation with higher doses of CNO may reveal a similar change in spontaneous dynamics. Alternatively, Gi-GPCR-induced [glu]_e_ may be dependent on visual stimuli and may play a role in regulating V1 circuit dynamics, such as through altering visual plasticity. This possibility could be assessed by combining astrocytic Gi-GPCR activation with a visual discrimination task. Further, performing these experiments in other primary sensory cortical areas may reveal generalized sensory mechanisms, while examining non-sensory cortical areas like frontal cortex could reveal more general astrocyte-neuron cortical circuit principles.

## Acknowledgements

The authors are grateful to members of the Poskanzer lab for helpful discussions. We thank Ilya Kolb, Jonathan Marvin, and Jeremy Hasseman for providing iGluSnFR and iGABASnFR2 viruses, and Jennifer Thompson for administrative support. K.P. is supported by NIH R01NS099254, NIH R01MH121446, NSF CAREER 1942360, and is a Chan Zuckerberg Biohub Investigator. T.V. was supported by the UCSF Genentech Fellowship.

## Methods

### Animals

All procedures were carried out in adult mice (C57Bl/6, P50–100) in accordance with protocols approved by the University of California, San Francisco Institutional Animal Care and Use Committee (IACUC). All animals were housed in a 12:12 light-dark cycle with food and water provided *ad libitum*. Male and female mice were used for all experiments. Following surgery, all animals were singly housed, to protect fiberoptic implants, with additional enrichment.

### Surgical procedures

Adult mice (C57Bl/6, P50–100) were administered dexamethasone (5mg/kg, s.c.) prior to surgery and anesthetized with isoflurane. For iGluSnFR experiments, two viruses, *AAV-GFAP*.*SF-iGluSnFR*.*AI184S* and *AAV5-GFAP-hM4D(Gi)-mCherry* were mixed in a 2:1 ratio and 600nL of the mixture was injected in V1 (−3.5mm, 2.5mm lateral from bregma). For iGABASnFR2 experiments, two viruses, *AAV-GFAP-iGABASnFR2* and *AAV5-GFAP-hM4D(Gi)-mCherry* were mixed in a 2:1 ratio and 600nL of the mixture was injected in V1 (−3.5mm, 2.5mm lateral from bregma). Injections were made 0.35mm from the pial surface followed by a 10-min wait for diffusion. Following viral injection, a fiberoptic cannula (Doric Lenses: 400μm inner core, 430μm outer core, 0.57 NA, 1.0mm length) was implanted 0.3mm below the pial surface and secured with a layer of superglue followed by C&B Metabond. Post-operative care included administration of 0.05mg/kg buprenorphine and 5mg/kg carpofen. Mice were given 2–3 weeks for recovery and viral expression.

### DREADD activation

Prior to fiber photometry recordings, mice were habituated to the recording setup (circular chamber lined with white ALPHA-dri bedding, and tethering via patchcords) for ∼30min. At the start of each experimental day, mice were weighed and connected to the recording apparatus. Immediately before the recording was started, CNO (1mg/kg) or saline (0.9%) was administered with an I.P. injection. CNO was diluted to 300μM in saline from a stock of 60mM each day, and the appropriate volume was measured for each mouse for a dose of 1mg/kg. An equal volume of saline was injected on control days. The sequence of CNO and saline control days was randomized amongst mice.

### Fiber photometry recording and preprocessing

Following injection of either CNO or saline, four recordings were made in the following order: spontaneous (no LED, 10-min), LED flash stimuli (10-min), spontaneous (no LED, 30-min), LED flash stimuli (10-min) for a total of 1-hour. A rig for fiber photometry recordings was used, similar to previously described^59,60^. Briefly, we used a Tucker-Davis Technologies RZ10X Processor with a Doric Lenses fluorescence mini-cube. A 473nm LED was used for the iGluSnFR and iGABASnFR2 excitation and a 405nm LED was used as an isobestic control. Both LEDs (Tucker-Davis Technologies, RZ10X Processor) went through a fluorescence mini-cube (Doric Lenses), and then through patchcords connected to a commutator to allow for free movement of the animal. After the commutator, a patchcord was connected to the fiber-optic cannula implanted in the animal. Fluorescence signals were reflected back through the mini-cube to a photoreceiver on the RZ10X Processor.

Raw fiber photometry data were minimally preprocessed in MATLAB, similar to previously reported^61^. First, the isobestic control channel was scaled to match the iGluSnFR or iGABASnFR2 channel. Next, the dF/F was calculated by subtracting the SnFR signal from the scaled isobestic channel and then dividing by the scaled isobestic channel. Lastly, the signal was detrended using the MATLAB function *detrend*.

### Visual stimuli

For visual stimulation, a blue LED light (ThorLabs, 470nm) was fed through a LED driver (ThorLabs), and connected through a patchcord attached above the circular chamber. The LED driver was controlled through the Tucker-Davis Technologies RZ10X processor to flash (illuminating the entire circular chamber), every 10–15s with a 500ms duration, throughout the fiber photometry recording. The timestamp of each LED onset was recorded alongside fiber photometry data for analysis.

### Analysis

#### Spontaneous dynamics

Peaks were automatically detected in the dF/F traces using the MATLAB function *peakfinder*, a noise-tolerant way to detect local maxima, and manually inspected for accuracy.

#### Stimulus-evoked dynamics

For each animal, the dF/F signal 3s before and 3s after each LED onset was averaged to obtain an event-triggered average (ETA) trace. Next, the timestamp for 5 events were identified within each ETA trace: (1) onset of early peak, (2) onset of late peak, (3) early peak, (4) late peak, and (5) trough. For the onsets of peaks (1 and 2), the derivative of each ETA trace was calculated and the negative-to-positive zero crossings were identified. For the early onset, the zero crossing that corresponded to the lowest value of the ETA trace 0–250ms from LED onset was chosen. For the late onset, the zero crossing that corresponded to the lowest value of the ETA trace 300–500ms from LED onset was chosen.

For the peak times (3 and 4), the derivative of each ETA trace was calculated and the positive-to-negative zero crossings were identified. For the early peak time, the zero crossing that corresponded to the maximum value of the ETA trace 200–400ms from LED onset was chosen. For the late peak time, the zero crossing that corresponded to the maximum value of the ETA trace 400–800ms from LED onset was chosen.

For the trough (5), the derivative of each ETA trace was calculated and the negative-to-positive zero crossings were identified. The zero crossing that corresponded to the lowest value of the ETA trace 500– 2800ms from LED onset was chosen as the trough time.

The decay of the early peak was identified by calculating the “decay amount”, by subtracting the maximum value from the minimum value between event 3 (“early” peak) and event 2 (“late” onset), and finding the time within this range when the ETA trace reached 33% the decay amount. The decay tau of the late peak was identified by fitting an exponential function to the ETA trace between event 4 (late peak) and event 5 (trough) using the MATLAB function *fit*.

### Immunohistochemistry

After physiology experiments were complete, mice were intracardially perfused with 4% PFA. Brains were collected, immersed in 4% PFA overnight at 4°C and switched to 30% sucrose for two days before being frozen on dry ice and stored at −80°C. Brains were sliced coronally (40μm thick) on a cryostat. Slices were stored in cryoprotectant at −20°C until staining. Slices were washed with 1x PBS, 5min x 3, then with 0.1% PBS-TX for 30min. Slices were next washed with 10% NGS (Invitrogen) for 1 hr, followed by an overnight incubation of 2% NGS, rat α-mCherry (1:2000, ThermoFischer), rabbit α-NeuN (1:1000, EMD Millipore), and chicken α-GFP (1:3000, Aves Lab) in 4°C. Slice were next rinsed with 1x PBS x 3 before incubating for 2 hr at room temperature with goat α-rat Alexa Fluor 555 (1:1000), goat α-rabbit 405 (1:1000), and goat α-chicken Alexa Fluor 488 (1:1000). Slices were washed again with PBS 3x for 5 min before slide-mounting and coverslipping using Fluoromount G.

To analyze colocalization of mCherry and NeuN at single-cell resolution, 63x images were taken on a spinning disk confocal (Zeiss). Slides were oil-immersed and three slices/animal (−3.8 and −2.3 from bregma) were imaged. In these slices, three ROIs were taken at random, spanning the total area in which virus was expressed. Colocalization of mCherry/GFP and NeuN was performed using Fiji’s Cell Counter Plugin.

### Quantification and statistical analysis

All statistical tests used, definition of center and dispersion measurements, and exact n values can be found for each figure in the corresponding figure legend. Additional information regarding statistical tests is described in the relevant sections. For all figures, significance levels are defined as the following: *: p<0.05, **: p<0.005, ***: p<0.0005.

## Data availability

All datasets are available on Dryad: doi:10.5061/dryad.h70rxwdmd

**Figure 2 –figure supplement 1:**
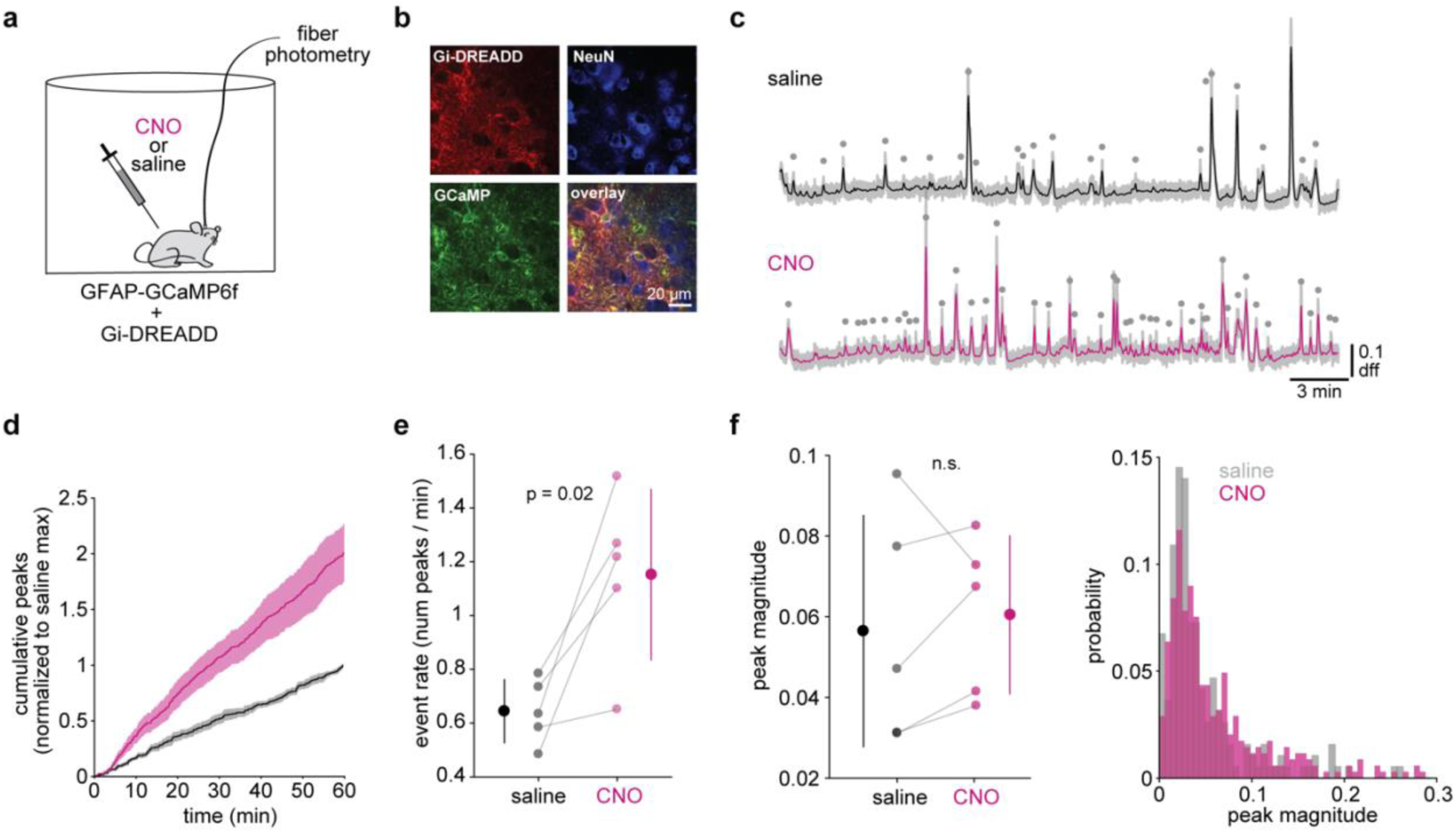
Gi-DREADD activation increases astrocyte Ca2+ events using fiber photometry. **(a)** Experimental paradigm. Photometry recordings of spontaneous astrocytic GCaMP activity were performed after I.P. injection of either CNO or saline. **(b)** Representative immunohistochemistry images of Gi-DREADDs, GCaMP, and NeuN. **(c)** Representative traces of spontaneous calcium activity recorded using fiber photometry. CNO injections (pink) resulted in more calcium transients (gray dots) compared to saline controls (black). **(d)** Cumulative Ca^2+^ event count for all mice after CNO (pink) shows higher event rate compared to saline (gray). Error bars = SEM. **(e)** Average event rate for each mouse was greater with CNO compared to saline (paired *t*-test). **(f)** The magnitude of Ca^2+^ events does not change with CNO administration, either in the distribution of all event magnitudes right) or in the average peak magnitude for each mouse (left). (For all panels, n = 9 mice. For panels e and f, paired t-test, error bars = SD)

